# Worms About Town: a citizen science outreach project discovers microsporidian parasites of nematodes through environmental sampling

**DOI:** 10.64898/2025.12.27.696690

**Authors:** Jonathan Tersigni, Edward B. James, Aaron W. Reinke

**Affiliations:** Department of Molecular Genetics, University of Toronto, Toronto, Ontario, Canada

**Keywords:** Community outreach, citizen science, nematodes, microsporidia, environmental sampling, infection, host-parasite systems

## Abstract

Environmental sampling of animal species can be a powerful tool for understanding aspects of host-parasite systems like species diversity and mechanisms of host specificity. To this end, we developed a citizen science project called Worms About Town to survey wild nematodes across Toronto, Ontario for microsporidia infection, which are natural fungal parasites of nematodes. Science communication and learning were emphasized through in-person outreach events and two-way discussion of project findings. Participants were equipped with tailored nematode collection kits to collect rotten plant substrates from local gardens and parks. In the first year of the project (2024), 13 participants acquired 127 samples, 60 of which contained nematodes and four were infected with microsporidia parasites. Microsporidia-infected nematodes were discovered from multiple types of plant substrates from various locations across Toronto. Sequencing of the microsporidia 18S rRNA gene of infected hosts identified two isolates of *Nematocida homosporus*, which is a known generalist microsporidian parasite of nematodes. The other parasites found in this study are undescribed species belonging to the *Pancytospora* and *Enteropsectra* genera, two microsporidia taxa that are distantly related to a clade of human-infecting microsporidia and known to infect wild nematodes. Three nematode isolates of the *Caenorhabditis* genus were discovered, including two isolates of *Caenorhabditis elegans*, a species not previously reported in Ontario. Both of the *N. homosporus* isolates were found in *C. elegans*, and one can infect a laboratory strain of *C. elegans*. The Worms About Town project will organize repeated wild nematode collections each year to determine natural factors that impact microsporidia infection in nematodes and to further characterize the diversity of this host-parasite system.

## INTRODUCTION

Microsporidia are obligate intracellular parasites capable of infecting many types of animals (Murareanu et al. 2021; Ruan et al. 2021). Microsporidia infection is typically detrimental for the host and impacts economically, ecologically, and agriculturally important animals including pollinators (Lang et al. 2023), farm animals (Aranguren Caro et al. 2021; Ruan et al. 2021; Moratal et al. 2023), and humans (Han et al. 2021). Although approximately 1700 species of microsporidia have been reported, this is likely a large underestimate of the true number of species (Franzen 2008; Larsen et al. 2017). Sampling efforts to identify novel microsporidian parasites have expanded our knowledge of both the host and phylogenetic diversity of microsporidia (Bojko et al. 2022; Willis and Reinke 2022).

Microsporidian biology is difficult to study as microsporidia are transcriptionally inactive while outside their hosts and exhibit extreme genome reduction with the smallest genomes of any known eukaryotes (Cuomo et al. 2012; Jespersen et al. 2022). Although microsporidia are thought to be as diverse as the species they infect and display different parasite lifestyles (Wadi and Reinke 2020), microsporidiosis is broadly treated via the same two drugs, fumagillin and albendazole, which exhibit host toxicity due to targeting essential and highly conserved eukaryotic proteins (Han and Weiss 2018). Therefore, there is a need to better understand the total diversity of microsporidia parasites and the interactions with their hosts to develop more effective and specific treatments.

Model organisms can be leveraged to study the enigmatic and recalcitrant microsporidian parasites. Microsporidia naturally infect zebrafish (Schuster et al. 2024), fruit flies, (Franzen et al. 2005), rodents, (Didier et al. 1995), and nematodes (Troemel et al. 2008) - all highly tractable and widely used model organisms in molecular biology. The nematode *Caenorhabditis elegans* and its natural microsporidian parasites have been used in screening experiments aiming to identify novel microsporidia-specific drug treatments (Murareanu et al. 2022; Huang et al. 2023) and to interrogate specific and general host responses to infection (Reddy et al. 2017; Willis et al. 2021). On a broader level, infection screens utilizing diverse nematode hosts and microsporidian parasites have identified distinct host specificity patterns and host tissue tropisms among nematode-infecting microsporidians. (Wadi et al. 2023); (Zhang et al. 2016)

Surveys for nematode-infecting microsporidians typically entail collection of wild nematodes from rotting plant substrates like fruits and vegetation, followed by parasite screening (Troemel et al. 2008); (Wadi et al. 2023); (Zhang et al. 2016); (Luallen et al. 2016); (van Himbeeck et al. 2025)). The methods and materials required to perform wild nematode isolation and parasite identification have been described (Tamim El Jarkass and Reinke 2024).

*Nematocida parisii* was the first microsporidian species identified that infects *C. elegans* (Troemel et al. 2008). Since then, collections of *Caenorhabditis* and other genera of nematodes have led to the identification of many *Nematocida* species, such as *N. ausubeli, N. major* (Zhang et al. 2016), *N. botruosus, N. cider, N. ferruginous* (Wadi et al. 2023), and *N. displodere* (Luallen et al. 2016). Other nematode-infecting genera have also been identified, including *Enteropsectra* and *Pancytospora* in *Oscheius* and *Caenorhabditis* nematodes, respectively (Zhang et al. 2016). These novel nematode-infecting microsporidia, unlike *Nematocida* species, retain RNA interference (RNAi) machinery in their genomes (Wadi and Reinke 2020) and may therefore be genetically tractable. Wild collections of nematodes and their microsporidian parasites have been instrumental in understanding the evolution and development of microsporidia (Wadi and Reinke 2020) and patterns of tissue tropism and host-specificity (Wadi et al. 2023); (Luallen et al. 2016); (Willis and Reinke 2022).

Inspired by previous community-aided projects to examine *Caenorhabditis* genetic diversity (Crombie et al. 2024) and to identify viruses of *C. elegans* (Ni and Sowa 2025), we sought to build a comprehensive wild nematode collection system while promoting scientific interest and outreach. To do this, we developed a citizen science project, called Worms About Town, that recruited local community members for surveying wild nematodes across Toronto, ON, Canada for microsporidia infections. This project aims to 1) discover, identify, and characterize nematode-infecting microsporidians; 2) monitor natural microsporidia infections in wild nematodes over time and determine environmental and evolutionary factors impacting microsporidia prevalence; and 3) build a network of citizen nematode-scientists throughout Toronto. These aims will be met by repeated annual collections of wild nematode samples across the city, and here we report the results of our first year of this project.

## RESULTS

### Environmental sampling by citizen scientists across Toronto discovered 60 wild nematode isolates and four microsporidia infections

To recruit community members for the collection of wild nematode samples, we pitched the Worms About Town project to local gardening groups and at science outreach events around Toronto (**Figure 1**). We demonstrated how to acquire samples from rotten plant substrates, seal them in petri dishes containing Nematode Growth Media (NGM), and screen petri dishes for nematodes using a light microscope (Tamim El Jarkass and Reinke 2024 - Basic Protocol 1) (**Supplemental Video 1**). Nematode sample collection kits were delivered to each project participant. These kits included NGM petri dishes, Ziplock bags, Parafilm strips, lab gloves, an infographic brochure (**Supplemental Figure S1**), and a sample collection guide/tracker (**Supplemental Figure S2**) to record the date, location, and plant substrate collected for each sample. Participants located rotten fruits and vegetation from their personal and/or local gardens/parks, tore off small pieces of the plant substrate, and placed them in NGM petri dishes. After ∼24 hours, the plant sample was removed from the dish. We then collected the petri dish samples and screened them for the presence of nematodes using a light microscope in the lab (**Figure 1**). Any nematodes discovered in the samples were then screened for microsporidia infection using the fluorescent dye Direct Yellow 96 (DY96), which stains chitin found in microsporidia spores (see methods; **Figure 1**).

**Figure 1:**
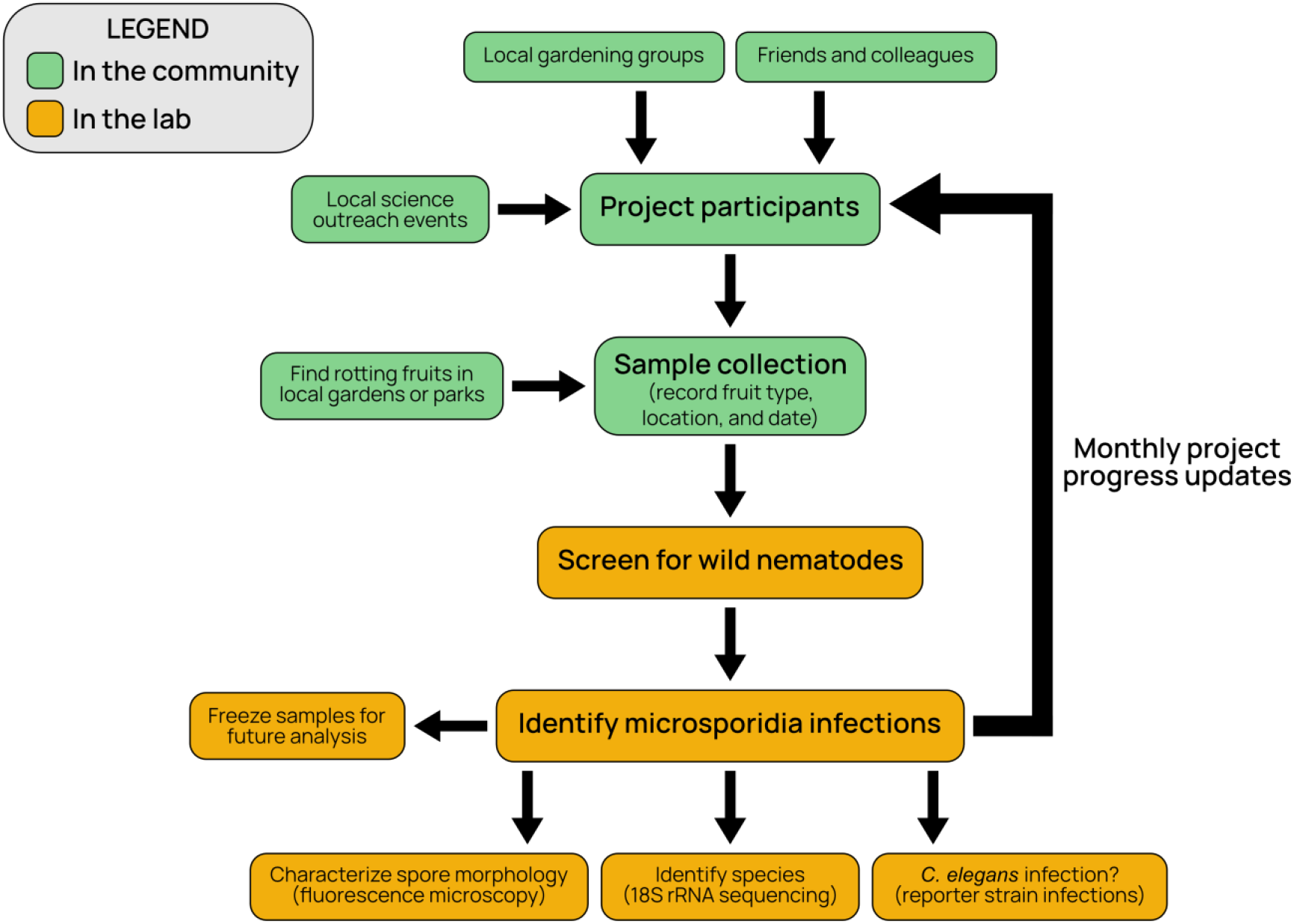
Flow chart outlining the Worms About Town project. Boxes in green indicate project steps involving the Toronto community (such as environmental sampling by participants), while boxes in orange indicate steps performed in the lab (such as screening samples for nematodes and microsporidia infections). Project updates and new findings were shared and discussed with all participants monthly.

During the inaugural year of the Worms About Town project, 13 participants collected 127 total samples across Toronto, ON, Canada from July to November of 2024 (**Figure 2A**). 60 (∼47%) samples contained nematodes, and four of the samples containing nematodes (∼7%) were infected with microsporidia spores (**Figure 2A**). Most samples were collected in September and October and the majority of nematode positive samples were found during these months.

**Figure 2:**
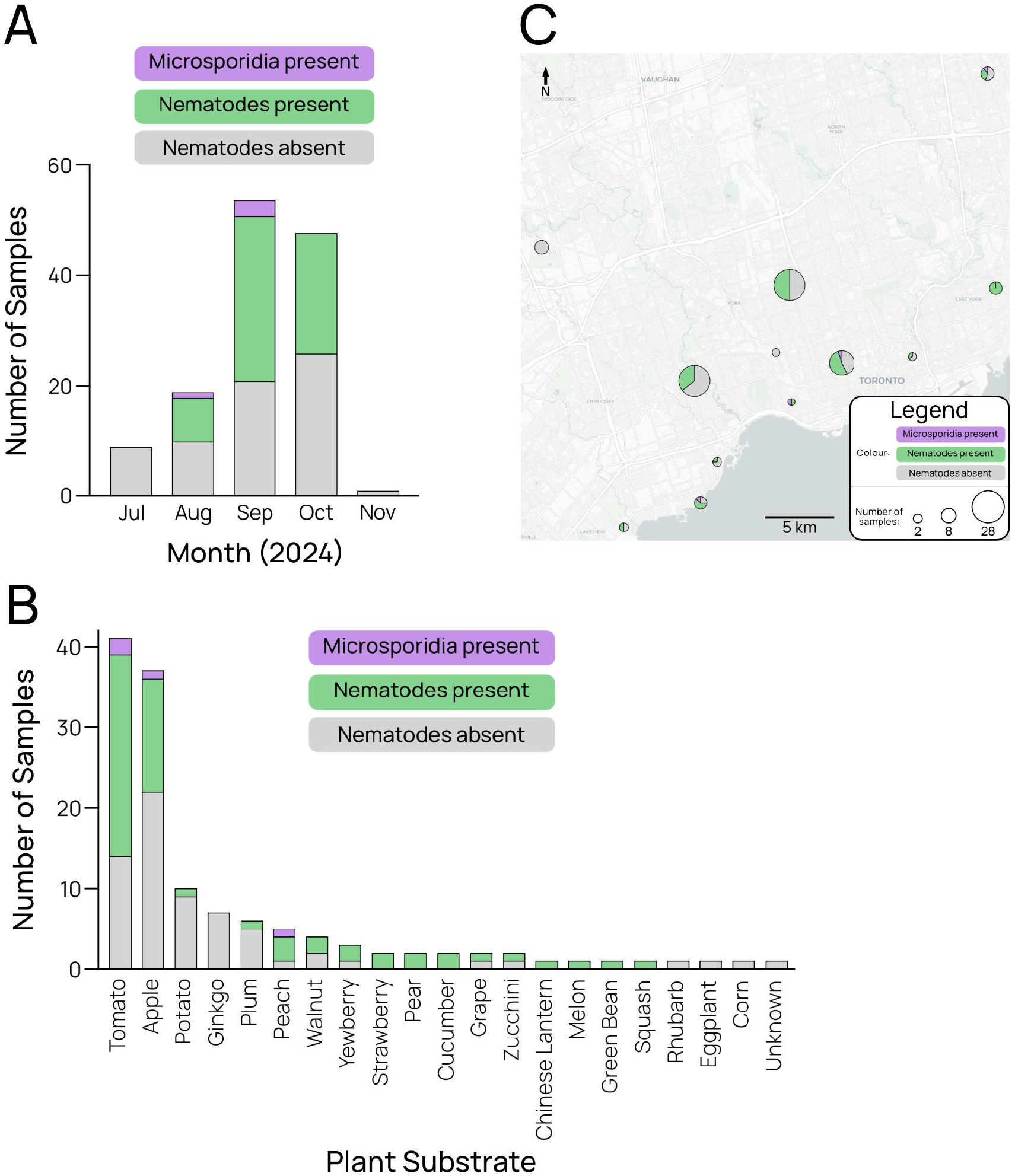
Community-driven sampling of wild nematodes and microsporidian parasites. The number of environmental samples collected in 2024 are organized by **A**) month, **B**) type of plant substrate, and **C**) location of collection across Toronto, Ontario. Grey indicates samples where no nematodes were found, while green and magenta indicate samples containing nematodes and microsporidia-infected nematodes, respectively. Pie chart size in **C**) indicates the number of samples collected at that location, while the black line indicates length of five kilometres. Map data is copyrighted by OpenStreetMap contributors and available from https://www.openstreetmap.org.

Samples collected in July and November did not contain nematodes, although these months had markedly fewer samples, nine and one respectively (**Figure 2A**), due to the low abundance of rotting plant substrates found in the environment. The 127 samples were comprised of 20 types of rotting plant substrates, the most prevalent being rotting tomato and apple, in which most nematodes were acquired (**Figure 2B**). Nematode-positive samples were found in 24 locations across Toronto, and all four microsporidia-positive samples came from different locations (**Figure 2C**). The success rates of finding wild nematodes and microsporidia infections in this study are similar to our lab’s previous findings and those reported in previous studies (Zhang et al. 2016); (van Himbeeck et al. 2025) and are a first point of comparison for the Worms About Town’s future surveys of Toronto.

### Characterization of four microsporidia-infected nematode isolates

We began characterization of the four microsporidia-containing nematode samples using fluorescent microscopy. The samples were named in a numerical fashion to identify the project participant and the number of samples they collected (for example, sample “8_13” indicates participant number 8, sample number 13). Each sample was stained with DY96 to determine spore morphology (**Figure 3A**). Approximate lengths and widths of microsporidia spores from each sample were measured (see methods) and determined to be 3.92 by 1.14 µm (3_3), 1.74 by 0.93 µm (15_2), 2.06 by 0.68 µm (8_13), and 2.20 by 0.93 µm (22_5) (**Figure 3A**).

**Figure 3:**
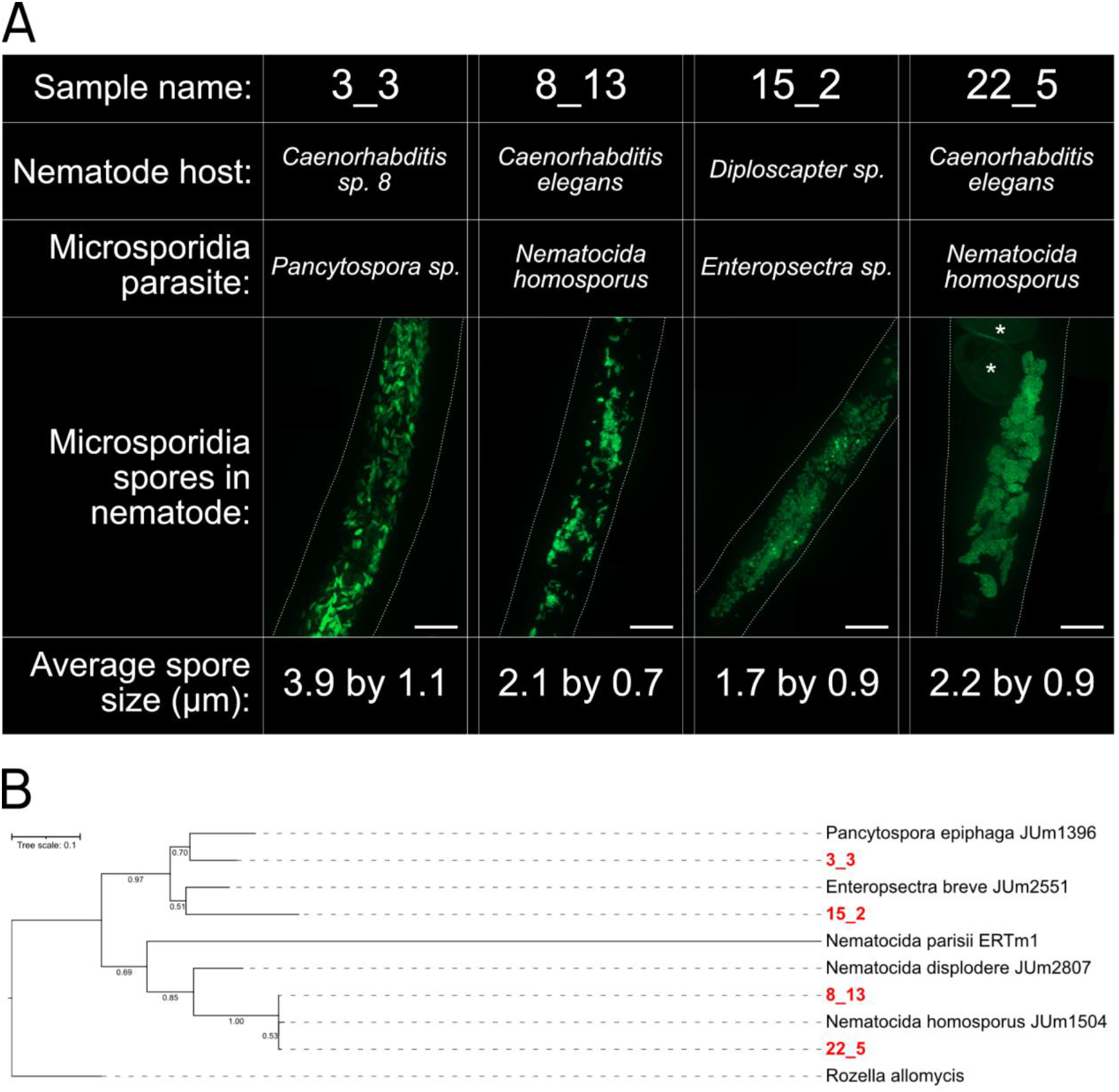
Four microsporidia-infected nematode samples were discovered and characterized. **A)** Representative fluorescent microscopy images of microsporidia spores (green) stained with DY96 within a wild-caught nematode (outlined with white dashes). DY96 also stains nematode embryos which are indicated by the asterisks in 22_5. The names of each collected sample are shown above each image and the average spore lengths and widths are shown below. The white scale bar indicates 20 µm in length. **B)** 18S rRNA phylogenetic tree of the four discovered microsporidia isolates, shown in red. Reference species, shown in black, are named followed by a reference strain identifier. The tree is rooted with the outgroup species *Rozella allomycis* and the bootstrap support for 1000 tree replicates which clustered together are shown at each node. Scale bar shows substitutions per site.

To molecularly identify the four wild nematode hosts and their microsporidia parasites found in this study, we isolated both host and parasite DNA by lysing infected nematodes (see methods). We then amplified and sequenced the nematode (Kiontke et al. 2004) and microsporidia 18S rRNA genes (Zhang et al. 2016) (Tamim El Jarkass and Reinke 2024) and aligned these sequences to previously published 18S sequences using BLAST (see methods). The identities of the nematode hosts and microsporidia parasites from each sample are displayed in **Figure 3A**. The microsporidia parasites in samples 3_3 and 15_2 were most closely related the microsporidia genera *Pancytospora* (87.91% nucleotide identity) and *Enteropsectra* (84.14% nucleotide identity), respectively, while both the 8_13 and 22_5 parasites were identified as *Nematocida homosporus* isolates (99.43% and 100% nucleotide identity, respectively). These identifications of 8_13 and 22_5 as isolates of *N. homosporus* using microsporidia 18S rRNA gene sequencing are generally consistent with the spore morphologies and sizes previously observed with our isolates being within 1-10% of the length and 4-23% of the width of the average of *N. homosporus* isolates (Zhang et al. 2016).

Nematodes from samples 3_3, 8_13, and 22_5 belong to the *Caenorhabditis* genus (100% nucleotide identity for all), while 15_2 nematodes belong to the *Diploscapter* genus (99.88% nucleotide identity). To determine the species of the *Caenorhabditis* nematode isolates, we amplified and sequenced the nematode ITS2 region (see methods), which can distinguish between species of *Caenorhabditis* (Kiontke et al. 2011). Aligning these sequences to previously published ones using BLAST determined that both the 8_13 and 22_5 nematodes are isolates of

*C. elegans* (100% nucleotide identity for both), while 3_3 nematodes are an isolate of *Caenorhabditis sp. 8* (100% nucleotide identity). To further verify that the 8_13 and 22_5 nematodes belong to the *C. elegans* species, we crossed them with males of a fluorescently labelled *C. elegans* strain (Au et al. 2019) (see methods). Fluorescent offspring were observed from both crosses (**Data S1**), indicating that successful mating occurred and demonstrating that 8_13 and 22_5 nematodes are isolates of *C. elegans*.

To gain an understanding of the relationships between the four newly discovered microsporidia and known microsporidia species, we constructed a phylogenetic tree based on their 18S rRNA sequences (**Figure 3B**) (see methods). The tree was rooted on *Rozella allomycis*, a relative of canonical microsporidia that is frequently used as an outgroup for creating microsporidia phylogenies. As expected, 3_3 and 15_2 microsporidia formed a clade with a *Pancytospora* isolate and *Enteropsectra* isolate, respectively, while both 8_13 and 22_5 microsporidia grouped together with a *Nematocida homosporus* isolate (**Figure 3B**).

### The *N. homosporus* isolate 22_5 can infect *C. elegans* in the laboratory

To determine if microsporidian parasites identified in this study can infect a laboratory strain of *C. elegans*, we utilized a previously published protocol for generating concentrated microsporidia spore solutions (Tamim El Jarkass and Reinke 2024). Spore solutions from wild samples are used to infect synchronized populations of the *C. elegans* strain ERT54, which is commonly used as a reporter strain for microsporidia infections in the laboratory (Bakowski et al. 2014), as well as the original host nematodes as a positive control for infection (**Figure 4A**). The *N. homosporus* isolate from sample 22_5 was found to successfully infect both *C. elegans* ERT54 and 22_5 (*C. elegans*) nematodes, as measured by quantifying the proportion of nematodes containing newly generated infective spores (**Figure 4B**) (see methods).

**Figure 4:**
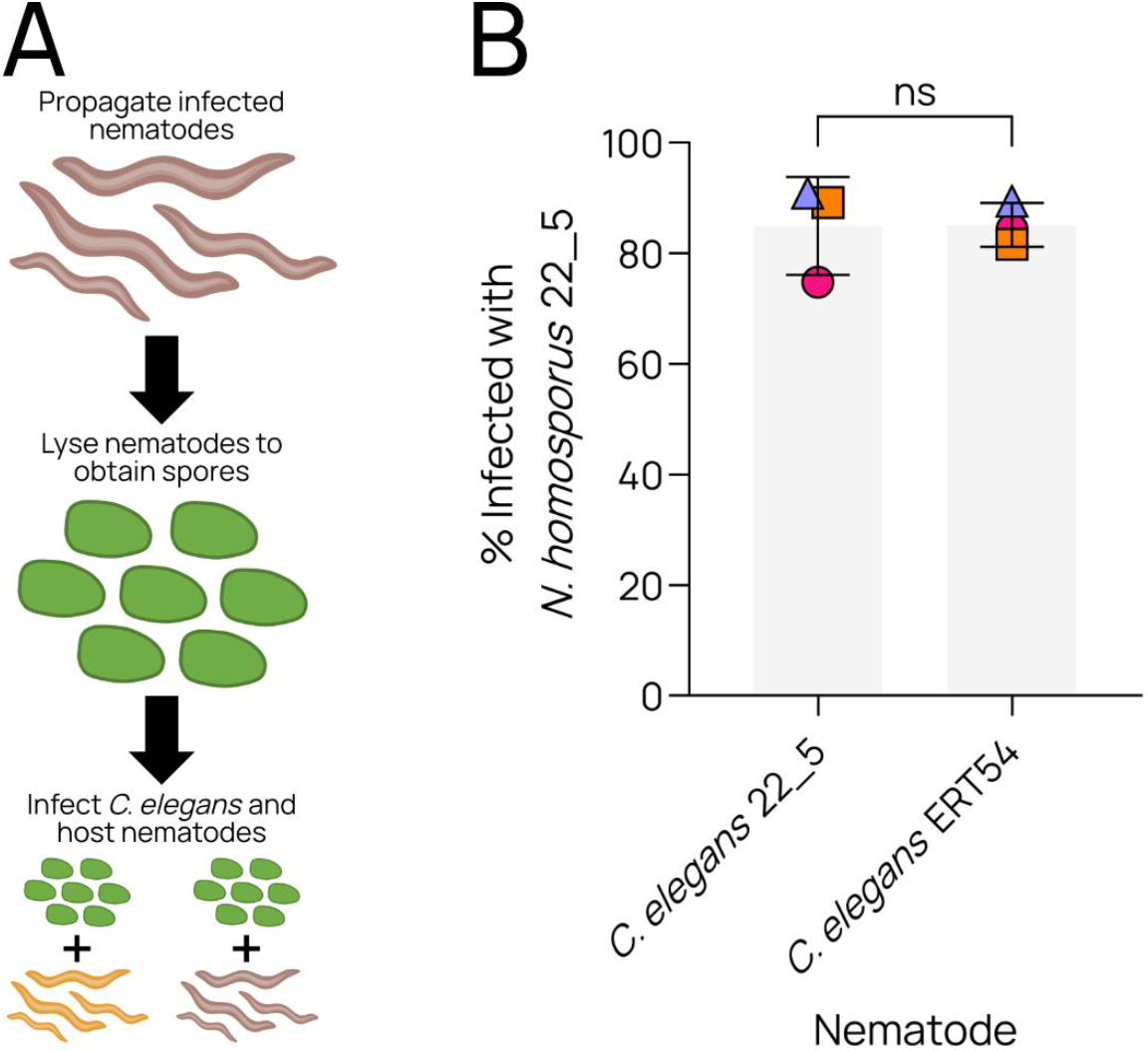
*N. homosporus* 22_5 infects a laboratory *C. elegans* strain. **A)** Schematic depicting how microsporidia from infected nematode samples were used to infect nematodes. Infected nematodes were grown on NGM plates, then lysed to obtain concentrated microsporidia spore solutions, which were used to infect either *C. elegans* or the original host nematodes as a positive control. **B)** The percent of nematodes (*C. elegans* ERT54 or *C. elegans* 22_5) containing newly produced *N. homosporus* 22_5 microsporidia spores after 96 hours post-infection. Individual replicates are indicated with different colours and shapes, while the mean of the three replicates is indicated with the bar. To determine statistical significance, a two-tailed paired t-test was performed on the data. “ns” = not significantly different.

## DISCUSSION

The first year of the Worms About Town science outreach project involved 13 members of the Toronto community. 127 environmental samples were collected, 60 samples contained wild nematodes, and four nematode samples were infected with microsporidia parasites. Two isolates of *Nematocida homosporus* were identified, a microsporidian species known for being a generalist parasite of multiple nematode hosts where it infects the intestinal tract. One of the *N. homosporus* microsporidia isolates, from sample 22_5, was demonstrated to be able to infect the genetic model *C. elegans* in a laboratory setting. Novel species of *Enteropsectra* and *Pancytospora* were also discovered. Species of these genera are particularly interesting as they are related to human-infecting microsporidia and unlike *Nematocida* microsporidia, retain their RNAi machinery (Wadi and Reinke 2020). Identifying additional isolates of these taxa that can infect *C. elegans* could present a unique opportunity to perform RNAi on microsporidia in the context of infection.

From our sampling, we expanded the geographic and host diversity of nematode-infecting microsporidia species. *N. homosporus* was previously reported to be found in *Oscheius tipulae* nematodes in France and *Rhabditella typhae* nematodes in Portugal (Zhang et al. 2016). *N. homosporus* was also previously found to infect *Caenorhabditis*, including *C. elegans*, in a laboratory setting (Zhang et al. 2016). Here we report the first identification of *N. homosporus* infecting *Caenorhabditis*, specifically *C. elegans*, in the wild, and also the first identification of this microsporidia species in Canada. Our collection also recorded the first microsporidian infection of a *Diploscapter* genus nematode – which belongs in a sister taxon to the *Caenorhabditis* genus (Kiontke and Fitch 2005), and two isolates of *C. elegans* nematodes. *C. elegans* are not known to be found in Ontario or surrounding regions, so our report of this species in Toronto suggests a possible range expansion (Frézal and Félix).

The Worms About Town project will continue to organize annual environmental sampling to 1) develop continued interest and involvement in microbiology and scientific research among members of the Toronto community; 2) determine the impact of natural factors like season, geography, and time on microsporidian infections of nematodes; 3) further our understanding of the natural prevalence and diversity of wild nematodes and their microsporidian parasites; and 4) uncover novel and/or interesting nematode-microsporidia combinations that could be utilized in the laboratory to provide insight into mechanisms of parasite host specificity and host susceptibility/resistance to infection.

## METHODS

### Environmental sampling by citizen scientists

We conducted a survey in the Greater Toronto Area throughout the summer and fall of 2024 for nematodes and their associated microsporidia. To allow a wide and consistent sampling area, we recruited citizen scientists at local science outreach events and via local gardening networks. We arranged to collect samples of rotting fruits and vegetation each month from late July to the beginning of November. To ensure samples were collected consistently and safely, we created an instruction/demonstration video and distributed protocol cards concisely describing our previously published wild nematode collection protocol (Tamim El Jarkass and Reinke 2024 - Basic Protocol 1). Briefly, volunteers collected small samples of rotten plant substrates (fruit and/or vegetation) from their garden or a local park and placed it on an *Escherichia coli* OP50-1 seeded 6 cm Nematode Growth Media (NGM) petri dish. Volunteers removed the plant substrate after 24 hours, whereupon we collected their petri dishes and screened for nematodes under a dissecting microscope. The type of plant substrate collected, the location of collection, and the date of collection for each sample were recorded by the participants.

### Sample nomenclature

Each project participant was assigned a unique reference number, and each of their samples were sequentially numbered. For example, the sample “22_5” is the fifth sample collected by our 22^nd^ participant. Some participants were assigned numbers and received collection kits without returning samples to us, hence the inflated participant number for 13 volunteers. Each of the four microsporidia-infected nematode isolates were assigned a designated lab strain name (AWR): 3_3 (AWR192), 8_13 (AWR193), 15_2 (AWR194), 22_5 (AWR195).

### Microsporidia spore staining and size measurements

To screen wild nematodes for microsporidia infection, we followed the protocol outlined in Tamim El Jarkass and Reinke (2024). Briefly, nematodes were washed off petri dishes with 1 mL of M9 into microcentrifuge tubes. Nematodes were pelleted by centrifuging at 1,400 rcf for 30 seconds and the supernatant was discarded. 1 mL of acetone was used to resuspend the nematode pellet and the nematodes were incubated at room temperature for 10 minutes to allow for fixation. After this period, the nematodes were again pelleted and the supernatant was discarded. The nematodes were washed twice with 1 mL of Phosphate Buffered Saline (PBS) supplemented with 0.1% Tween-20 detergent. The nematodes were then resuspended with 500 µL of DY96 staining solution (1X PBS, 0.1% Tween-20, 0.1% SDS, 0.02 mg/mL Direct Yellow 96) and incubated on a tube rotator for 30 minutes covered in aluminium foil to protect from light. The nematodes were then pelleted and resuspended with 16 µL of Everbrite Mounting Medium, which was then added to a microscope slide and covered with a coverslip. The DY96-stained nematodes were imaged on a ZEISS Axio Imager 2 at magnifications ranging from 5X to 63X. To quantify spore sizes within nematodes, a z-stack image was captured using the 63X magnification and an apotome maximum intensity projection image was created. The lengths and widths of at least 20 spores from each sample were measured on these images using the line tool on ImageJ. Freezer stock solutions of each microsporidia-positive sample were created by washing two 10 cm plates of nematodes with 2 mL of Freezing medium (see Tamim El Jarkass and Reinke 2024 - Basic Protocol 2) and stored at –80 °C to allow for future analysis and comparison with future collections.

### 18S rRNA gene sequencing and phylogenetic analysis

To identify the nematodes and microsporidia in the four infected samples, we adapted the protocol outlined in Tamim El Jarkass and Reinke (2024). Briefly, nematodes from the four microsporidia-infected samples were continuously propagated on 10 cm NGM petri dishes seeded with *E. coli* OP50-1. To generate DNA template for 18S rRNA PCR amplification, nematodes from two 10 cm petri dishes per infected sample were washed into microcentrifuge tubes using 3 mL of M9. The nematodes were then washed three times with M9 supplemented with 0.1% Tween-20 to remove contaminating microbes. 25 µL of the final nematode pellet was combined with 25 µL of single worm lysis buffer (see Tamim El Jarkass and Reinke 2024 - Basic Protocol 3) and incubated at 65 °C for 60 minutes then 95 °C for 15 minutes in a thermocycler to lyse nematodes and microsporidia spores. The resulting lysate contains both nematode and microsporidia DNA and was used as template for PCR.

The forward primer V1F (CACCAGGTTGATTCTGCCTGAC) and reverse primer 18SR1492 (GGAAACCTTGTTACGACTT) (Zhang et al. 2016) were used to amplify 22_5 and 15_2 microsporidia, while V1F and the reverse primer Micuni3R (ATTACCGCGGMTGCTGGCAC) (Doliwa et al. 2023) amplified 3_3 and 8_13 microsporidia. To amplify nematode 18S rRNA, the universal nematode forward primer G18S4a (GCTCAAGTAAAAGATTAAGCCATGC) and reverse primer DF18S-8 (GTTTACGGTCAGAACTASGGCGG) were used (Kiontke et al. 2004). To amplify the *Caenorhabditis* ITS2 region, the forward primer 5.8S-1 (CTGCGTTACTTACCACGAATTGCARAC) and reverse primer KK28S-4 (GCGGTATTTGCTACTACCAYYAMGATCTGC) were used (Kiontke et al. 2011). Sanger-sequencing of the PCR products was performed and regions of poor quality within each sequence were trimmed to generate a high-quality sequence. These sequences (NCBI accessions W, X, Y, and Z) were used as queries for nucleotide BLAST searches to ascertain nematode and microsporidia species identities (Tamim El Jarkass and Reinke 2024 - Basic Protocol 3).

A maximum-likelihood phylogeny was generated using our sequenced 18S rRNA genes and reference microsporidia 18S rRNA sequences from previous studies (Zhang et al. 2016). Sequences were trimmed to exclude ambiguous bases before alignment via MUSCLE (Edgar 2004), and tree construction was done using MEGA12 (Kumar et al. 2024). Model selection via MEGA12 suggested the Tamura model of nucleotide substitution (Tamura 1992) with 22.59% of sites treated as evolutionarily invariable (‘T92 + I’ model). We used a partial deletion cut-off of 80% to focus on near-ubiquitous sequences, reducing our dataset to 335 positions, and bootstrapped 1,000 times. The tree was then manually rooted on the microsporidia outgroup *Rozella allomycis*.

### Wild nematode mating with a *C. elegans* GFP reporter strain

To confirm that the 8_13 and 22_5 wild nematodes are *C. elegans* isolates, we mated them with the *C. elegans* strain RG3405, which contains a GFP reporter (Au et al. 2019). As a negative control, we also included 3_3 nematodes, which from our 18S rRNA analysis were found to belong to the *Caenorhabditis* genus (*Caenorhabditis sp. 8*) but not *C. elegans* (see results). Three larval stage 4 (L4) hermaphrodites/females of each wild nematode isolate and five L4 RG3405 males were placed on an individual 6 cm NGM petri dish seeded with *E. coli* OP50-1 for 24 hours for mating. After the mating period, each of the three hermaphrodites/females were transferred to a fresh 6 cm NGM OP50-1 dish. Two days after this transfer, the number of GFP-positive and GFP-negative fluorescence offspring were counted on the dish. Over 100 progenies were quantified per dish, where possible (**Data S1**).

### Infection of *C. elegans* by environmental microsporidia

To generate concentrated microsporidia spore solutions of microsporidia parasites from each infected nematode sample, nematodes from three densely populated 10 cm NGM petri dishes (approximately 2,500 nematodes per dish) for each sample were resuspended with 2 mL of sterile Millipore water into a 2 mL microcentrifuge tube. The tubes were spun at 1400 rcf for one minute to pellet nematodes, the supernatant was discarded, and the nematode pellets were resuspended with 1 mL of water. Two more washes were performed to dilute contaminating microbes. A final 1 mL volume of water was used to resuspend the nematode pellets, to which approximately 500 µL of 2 mm zirconia silica disruption beads were added. The tubes were vortexed in a bead disruptor at 5000 rpm for 3 minutes to lyse nematodes. The lysate was passed through a 5.0 µm sterile syringe filter to remove debris from the spore solution. The filter was then washed with 0.5 mL of water to gather spores remaining on the filter. The filtered spore solution was aliquoted into 50 µL volumes in 1.7 mL microcentrifuge tubes and stored at –80 °C.

To determine if the wild microsporidia isolates could infect a lab strain of *C. elegans*, 50 µL of spore solution derived from each sample was added to 350 µL of sterile M9. These solutions were then used to resuspend a pellet of 1200 bleach-synchronized L1s of *C. elegans* ERT54 or one of the four wild nematode hosts. The mixtures were individually plated on 6 cm *E. coli* OP50-1 seeded NGM petri dishes, dried in a clean cabinet, and stored at 21 °C. 96 hours after plating, nematodes were washed off the dishes with M9, fixed with acetone, and stained with DY96 as previously described to discern newly produced microsporidia spores.

## ACKNOWLEDGMENTS

We would like to thank all current and past members of the Worms About Town project who contributed their time and effort to sample wild nematodes. Project participants from the first year of Worms About Town who consented to the inclusion of their names are: Joseph Durand, Helen Elliot, Leslie Ferguson, Alissa Hamilton, and Tony Likhachov. We would also like to thank Reinke lab members Winnie Zhao and Meng A. Xiao for assisting in delivering nematode collection kits. This work was supported by a Canadian Institutes of Health Research grant no. 400784 (to A. W. R.). E. B. J. was supported by an Emerging & Pandemic Infections Consortium (EPIC) Convergence Fellowship and J. T. was supported by an Ontario Graduate Scholarship, EPIC Doctoral Research Award, and Natural Science & Engineering Research Council (NSERC) Post-graduate Scholarship.

## AUTHOR CONTRIBUTIONS

**Jonathan Tersigni & Edward B. James:** conceptualization, data collection and analysis, manuscript drafting.

**Aaron W. Reinke**: conceptualization, data analysis, manuscript drafting.

## DATA AVAILABILITY

Microsporidia 18S rRNA sequences are available under the GenBank accession numbers PX741243 (3_3), PX741244 (8_13), PX741245 (15_2), and PX741246 (22_5). Nematode 18S rRNA and *Caenorhabditis* ITS2 sequences are available under the GenBank accession numbers PX741247 and PX741251 (3_3), PX741248 and PX741252 (8_13), PX741249 (15_2), and PX741250 and PX741253 (22_5), respectively. All data from the study are available in **Data S1**.

## SUPPLEMENTAL MATERIAL

**Supplemental Figure S1:**
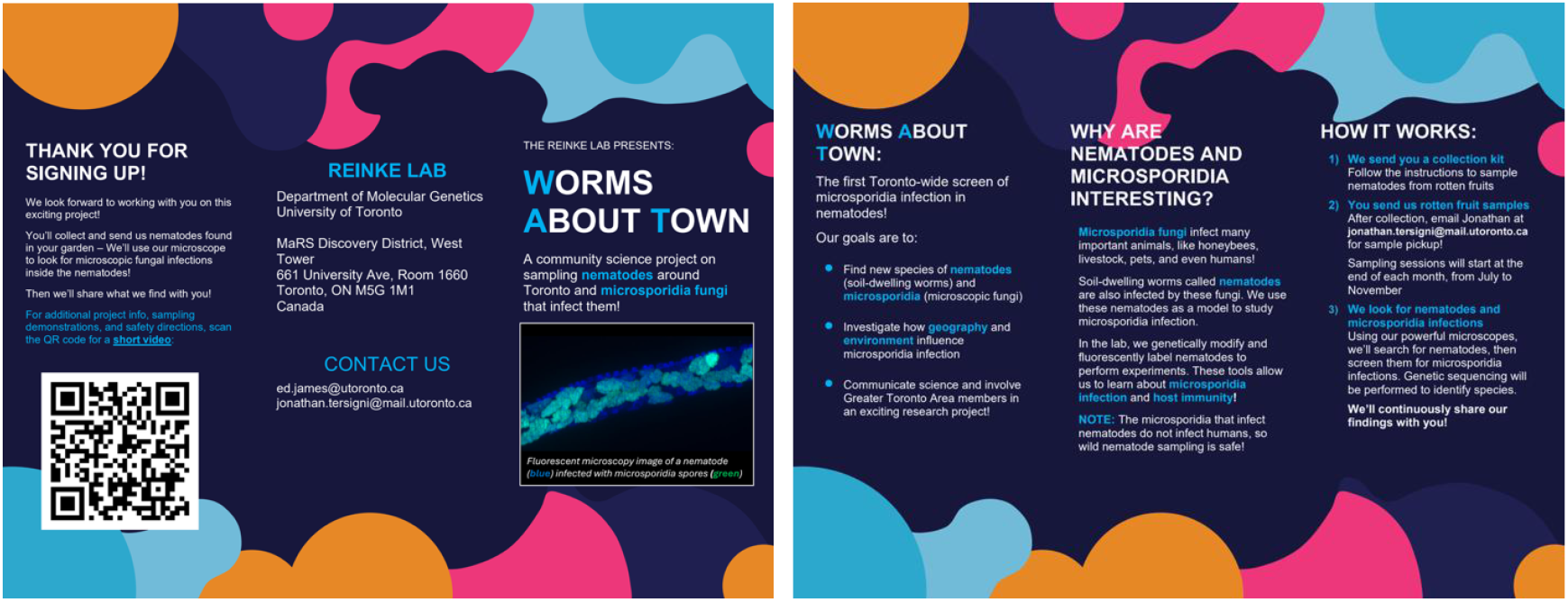
Trifold brochure detailing the Worms About Town project. This document was printed double-sided and included in the nematode collection kit distributed to all project participants. This document is available on the Reinke lab website (https://www.reinkelab.org/worms-about-town)

**Supplemental Figure S2:**
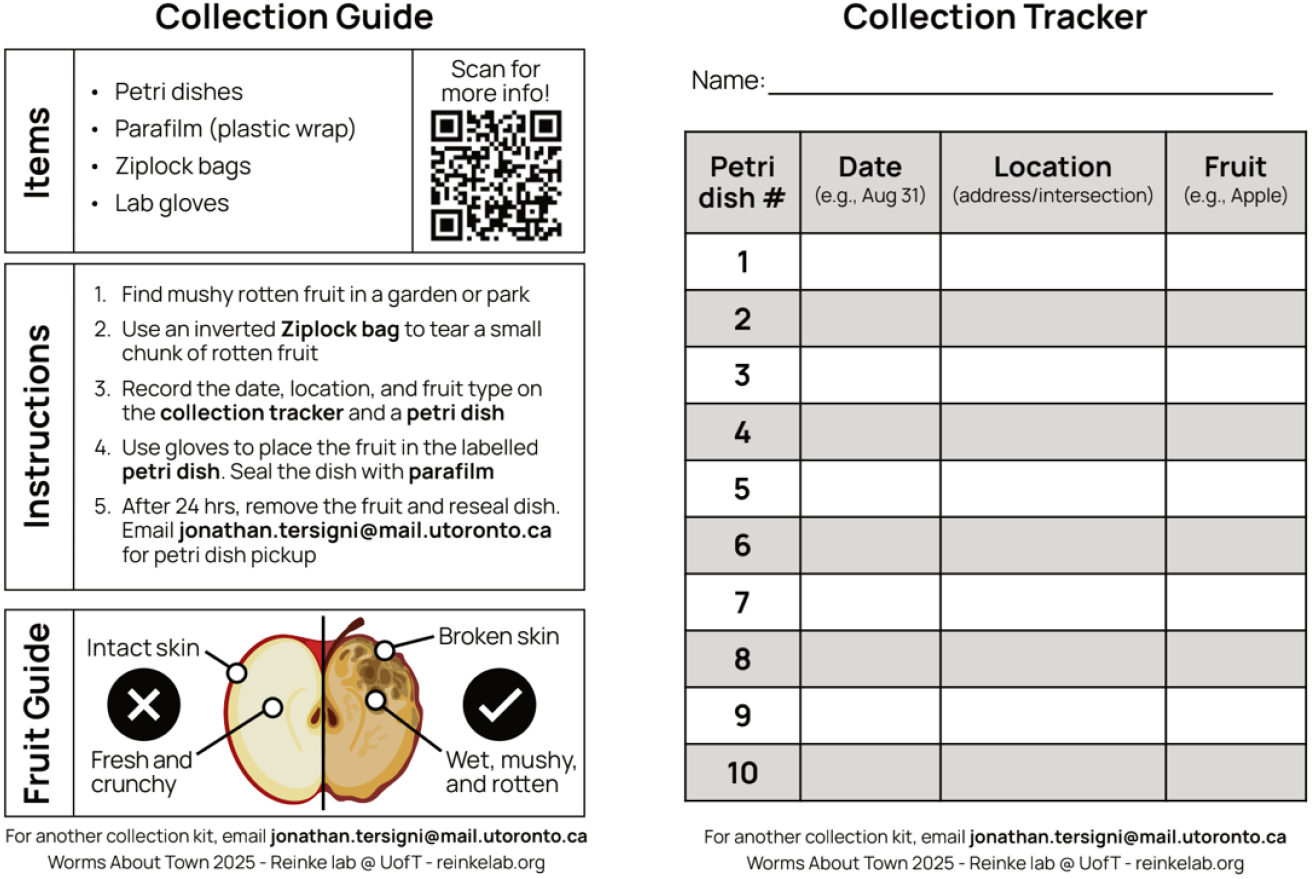
Nematode sample collection guide and tracker. This document was printed double-sided and included in the nematode collection kit distributed to all project participants. Participants recorded information for 10 collected samples in the Collection Tracker sheet, which was returned during sample pick-up. This document is available on the Reinke lab website (https://www.reinkelab.org/worms-about-town)

**Supplemental Video 1: Instructional video that explains the Worms About Town project and demonstrates how to collect environmental nematode samples from rotten fruit and vegetation**. This video is available on the Reinke lab website (https://www.reinkelab.org/worms-about-town) and YouTube (https://youtu.be/KmfU8mXRLJw?si=qIti8RUNMAz3wL).

